# Amplification, not spreading limits rate of tau aggregate accumulation in Alzheimer’s disease

**DOI:** 10.1101/2020.11.16.384727

**Authors:** Georg Meisl, Yukun Zuo, Kieren Allinson, Timothy Rittman, Sarah DeVos, Justin S. Sanchez, Catherine K Xu, Karen E Duff, Keith A. Johnson, James B Rowe, Bradley T Hyman, Tuomas P J Knowles, David Klenerman

## Abstract

Both the replication of protein aggregates and their spreading throughout the brain are implicated in the progression of Alzheimer’s disease (AD). However, the rates of these processes are unknown and the identity of the rate-determining process in humans has therefore remained elusive. By bringing together chemical kinetics with measurements of tau seeds and aggregates across brain regions, we are able to quantify their replication rate in human brains. Remarkably, we obtain comparable rates in several different datasets, with 5 different methods of tau quantification, from seed amplification assays *in vitro* to tau PET studies in living patients. Our results suggest that the overall rate of accumulation of tau in neocortical regions is limited not by spreading between brain regions but by local replication, which doubles the number of seeds every ~5 years. Thus, we propose that limiting local replication constitutes the most promising strategy to control tau accumulation during AD.

## Introduction

Alzheimer’s disease (AD), similar to several other aggregation-associated neuro-degenerative diseases (1, 2), is characterized by a progressive decline in health over the course of several years, with symptoms often only becoming apparent years after onset of pathological changes in the brain. The processes that are believed to be of critical importance in the development of AD are the aggregation of the initially soluble Aβ into plaques and tau proteins into neurofibrillary tangles (3). While Aβ aggregation is believed to be an important event, clinical symptoms, atrophy and brain damage correlate best with the appearance of tau aggregates (4). Tau aggregates have the ability to self-replicate and such replication competent aggregates are referred to as proteopathic seeds. Once an initial seed is present it can replicate to form a large number of new seeds. Indeed, synthetic tau filaments made from recombinant protein as well as filamentous material extracted from tau mouse models or AD brains have been shown to act as seeds in various model systems and initiate tau pathology (5–9). Furthermore, several mouse model systems provide evidence that seeds spread from the regions in which they are initially formed to other regions of the brain and trigger aggregation there (5, 10–12). The molecular processes that lead to tau seed replication and spreading are not known in detail, but based on animal models, are postulated to involve aggregation, transport down axons, release, uptake and finally replication in the recipient neuron.

It has been suggested that the patterns of location and abundance of tau neurofibrillary tangles observed in post-mortem AD brains, which form the basis for the classification of AD into Braak stages (13,14), arise from the spread of tau seeds along well-established connections through the brain. If the rate of this spread is slow enough (8,12), and assuming that aggregation begins in a single location, spreading from one brain region to the next could be the limiting factor in disease progression (7, 15). This idea underlies several current clinical trials with, for example, anti-tau antibodies.

However, while both the replication and spatial spreading of seeds occur *in vivo*, a key unanswered question in the study of AD in particular, and aggregation-related diseases in general, is whether the replication of seeds or their spread over longer length-scales, between brain regions, determines their kinetics over the time-scales of human disease, and at what rate this process occurs.

In this work we establish a general theoretical framework to determine the rates governing tau accumulation and apply it to measurements of seed concentrations in AD brains to determine the rate-limiting process and calculate the associated reaction rates. Our model is formulated in terms of general classes of processes and is therefore able to describe the wide range of possible mechanisms of replication and spreading *in vivo* and determine the effective rates of these processes. This fundamental model naturally results in two limits, where either long range spreading or local replication dominate the kinetics of tau accumulation.

To allow precise statements to be made based on our models, we give here clear definitions of a number of terms as used throughout this work (see also Fig. 1a,b): *Replication* is the process by which one seed can grow and multiply to become two (or more) seeds which are both capable of further replication. Its subprocesses naturally fall into two categories, growth and multiplication. *Growth* processes increase the size of a given seed, by the addition of more proteins. *Multiplication* processes increase the number of growth competent seeds and encompass a wide range of processes, from simple fragmentation of seeds, to indirectly induced multiplication via other biological processes. Growth and multiplication couple together, as new seeds have to mature by growth before multiplying again. *Aggregate* refers to all aggregated species. *Seeds* are species that are replication competent. *Spreading* is any process that results in the spatial relocation of aggregates, within a cell or from cell to cell, including diffusion as well as any active transport processes.

**Figure 1:**
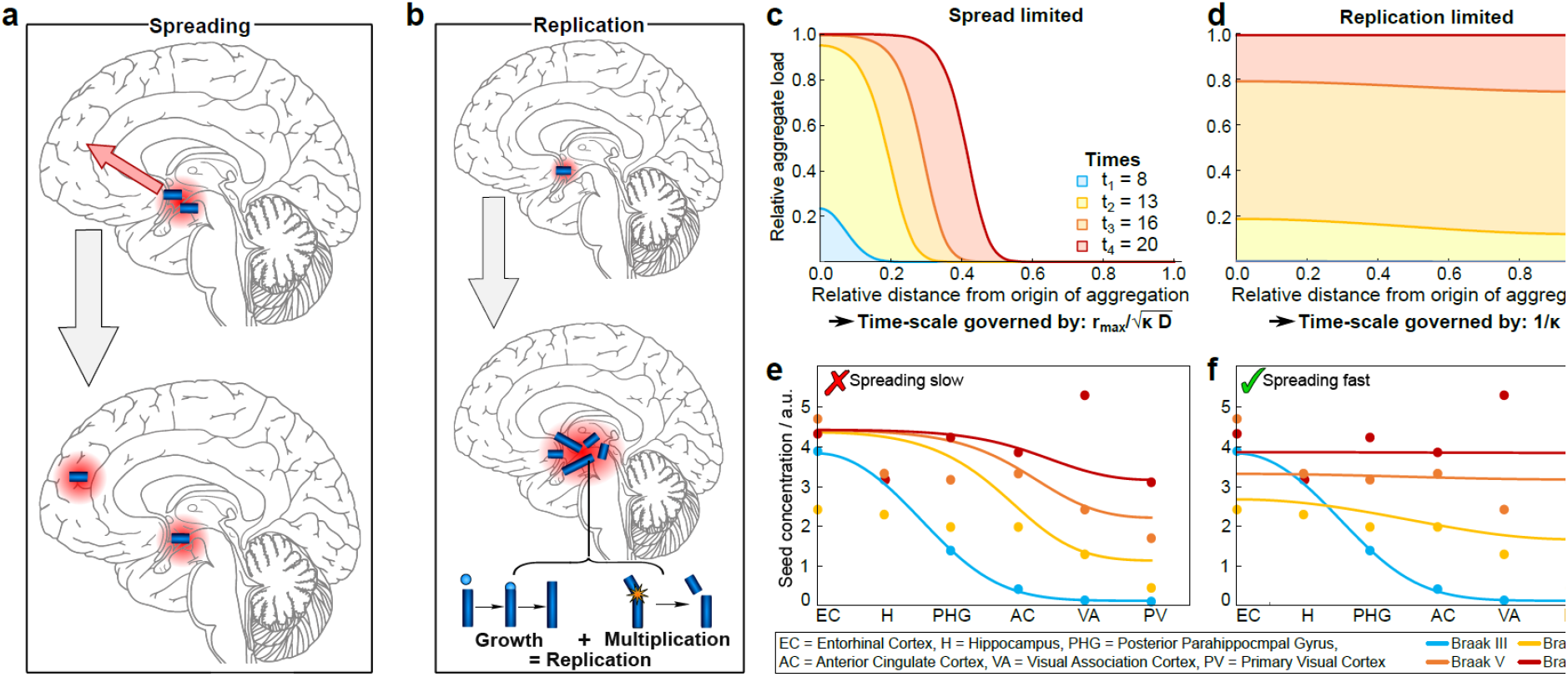
Illustration of the key processes in tau aggregate formation. **a** Spreading denotes spatial relocation of an existing aggregate. **b** Replication, the overall process that leads from one seed to many, is composed of subprocesses that naturally fall into two categories, those that increase seed number (multiplication) and those that increase the size of a given seed (growth). **c-d** Seed load as a function of time and distance, from the solution of equation 1, in the two limits. In c spread is slow (*D* = 0.00025 (unit length)^2^ years^−1^) and the system is spreading-limited, in d spread is fast (*D* = 0:025 (unit length)^2^ years^−1^), thus the system is replication-limited (see methods for an approximate conversion from these reduced units). **e** Dots show the experimentally measured distribution of tau seeds, sampled in several brain regions, at different stages of the disease (16). The regions from left to right correspond to increasing distance from the entorhinal cortex where aggregates first appear. Dashed lines are a guide to the eye. Solid lines are the results of a simulation of the data from Braak stage III onwards, assuming that long-range spreading is slow. **f** As in e, but the solid lines are the results of a simulation in which spreading is fast.

## Results

Mathematical models predict two limiting behaviours. We develop a general model by considering the different fundamental classes of processes and grouping together similar phenomena into one effective term. The level of coarse-graining is dictated by the detail of the experimental data available, ensuring that the simplest model consistent with the data is used. The fundamental model that includes these processes takes the form of a spatially-dependent reaction equation (17, 18):

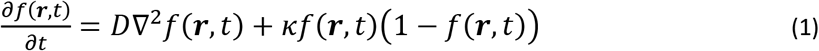

where *f(***r**, *t*) is the seed concentration relative to some maximal concentration, *P*_max_, at time *t* and position **r**. *D* is an effective diffusion coefficient that determines the speed of spreading and *κ* is an effective replication rate. The first term on the right-hand side of equation 1 accounts for spreading, in the form of an effective diffusion term. This common mathematical description of transport covers a wide range of active and passive processes in addition to simle diffusion (19). The second term accounts for replication, producing an initially exponential increase in seeds, which levels off as the maximal seed concentration is approached. The introduction of this maximal concentration is motivated by basic physical considerations, as well as experimental observations of such a limit.

If we initiate the reaction by introducing a small concentration of seeds at one location, two limiting regimes naturally emerge from the description in equation 1, which we refer to as the replication-limited and the spreading-limited cases, see Fig. 1 (for mathematical details see SI section 2.1). In a replication-limited regime, the overall timescale of the reaction is determined exclusively by the replication of seeds, which is the case when significant amounts of seeds are present throughout the reaction volume before the limit in seed concentration is reached anywhere (see Fig. 1d). This can be achieved both by the fast spreading of an initially localised distribution or alternatively by the initial presence of a small concentration of seeds everywhere throughout the reaction volume. Experimental realisations of this latter scenario would be systems in which seeds are introduced globally, or systems in which the spontaneous formation of seeds, directly from monomers, happens at many locations throughout the brain. By contrast, in a spreading-limited regime, replication is so fast that each region reaches the maximal seed concentration before a significant amount of seeds can spread to the next region. At all times most regions are either essentially free of seeds or at the limiting concentration of seeds. There is a clear propagation front moving through the reaction volume (see Fig. 1c).

The rates of spreading and the rates of replication as well as the size of the system and, crucially, the initial distribution of seeds determine which limit the system is in. These limiting regimes are a general feature and emerge regardless of whether spread is assumed to proceed directly through 3-dimensional space, or along specific axonal pathways (see SI sections 2.2 and 2.3).

### Behaviour of tau seeds in human AD is limited by local replication at later stages

Utilising the ability of tau seeds to replicate, low concentrations of these replication competent aggregates can be detected in brain samples using an amplification assay. In DeVos *et al.* (16) we measured the seed activity in 6 different brain regions from 29 individuals with AD neuropathological changes from Braak II to Braak VI, and found that regions that contained NFT, as well as areas that were presumed to contain synaptic projections of brain areas that had NFT, and found detectable seeding activity one or two synapses away from areas that had developed NFT. From these data, together with further measurements by the same technique from Furman *et al.* (20) and Kaufman *et al.* (21), as well as measurements by orthogonal methods using neuropathological approaches in human disease from Gomez-Isla *et al.* (22) and longitudinal *in vivo* tau PET imaging, we here determine the rates and rate-determining steps of tau accumulation during AD.

The spatial distribution of seeds at later Braak stages is shown in Fig. 1e. Three key observations guide the further analysis of these data: (i) The seed concentrations appear to increase in a concerted manner in the neocortical brain regions. (ii) Spatial inhomogeneity is most pronounced in the early Braak stages, up to stage III (see SI Fig. 5). (iii) There is a low but significant concentration of seeds even in the neocortical regions already prior to Braak stage III. The later stage behaviour appears qualitatively distinct from that of the very early Braak stages during which the majority of seeds is found in the Entorhinal Cortex (EC), Hippocampus and Posterior Parahippocampal Gyrus (PHG). During these early stages, high seed concentrations appear to be confined to the EC and Hippocampus and the concentrations in all other regions remain relatively low. This slow growing state, prior in many cases to the deposition of amyloid in the neocortex and the initiation of AD-like symptoms, appears to be relatively stable compared to the rate of doubling in later Braak stages where amyloid is also deposited. For the immediate model, we focus on the faster, disease associated phase of the disease. We simulate the situation by imposing the following initial condition in equation 1: at time 0 the concentration of seeds is modelled on the distribution measured in Braak stage III. The measured brain regions are represented by equally spaced locations in our simulations. This choice of effective distance reflects the fact that the number of synaptic connections to be crossed to go from one region to the region associated with the next higher Braak stage is similar for all regions. To obtain a measure for time, we convert from Braak stage to the length of time for which each stage lasts based on the extensive dataset by Braak *et al.* (14) (details see Materials and Methods, Fig. 4) (23). As the switch between the relatively constant distribution of the initial stages, and the global increase of the later stages happens between Braak stage III and VI, we choose our time 0 to lie between Braak stage III and VI. This value is optimised, but the time between consecutive Braak stages is fixed and the replication rate is also fixed to that obtained in the fits shown in Fig. 2.

**Figure 2:**
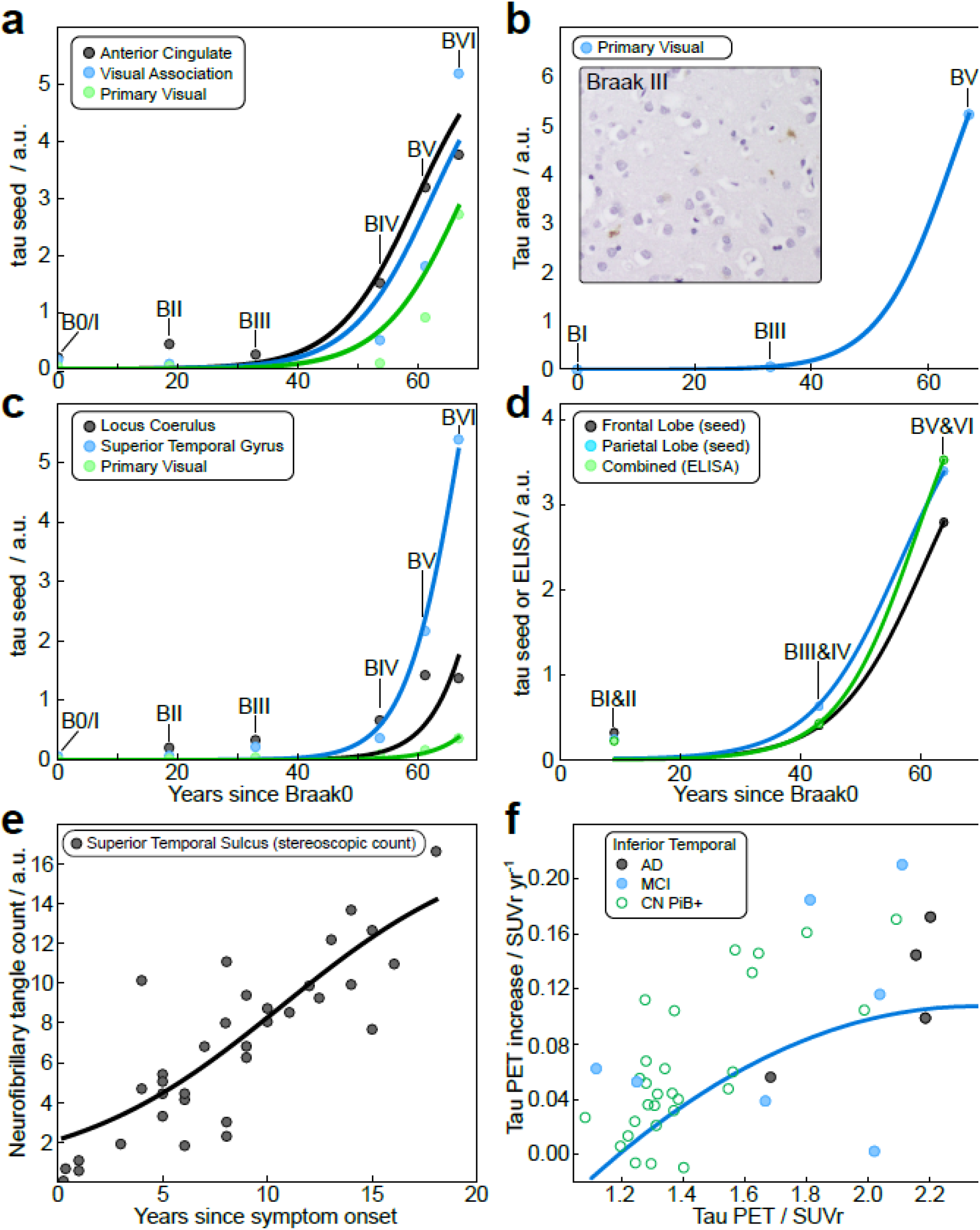
To determine the replication rate *κ*, the solution to equation 1 in the replication limit was fitted to **a** the temporal evolution of seed concentration in the neocortical regions, **b** the aggregate amounts measured by AT8 staining of brain slices from the primary visual cortex, (Images at Braak stages 0/I, III and VI were thresholded for AT8 response quantification, normalised by the number of cells) Inset: example image at Braak stage III. **c-d** to seed measurements from Kaufman *et al.* in c and seed and ELISA measurements from Furman *et al.* (20) in d and **e** stereological counts of neurofibrillary tangles from Gomez-Isla *et al.* (22). **f** shows longitudinal tau PET (^18^F-Flortaucipir) measurements form Sanchez *et al.* (24) which determined the rate of change in tau signal over consecutive measurements in the same patient. The data are shown as rate of change against total signal, the solid line is a prediction from analysis of the AD and MCI (mild cognitive impairment) patients. Also shown, but not included in the analysis, are cognitively normal patients with a positive amyloid beta PET signal (CN PiB+).

**Figure 4:**
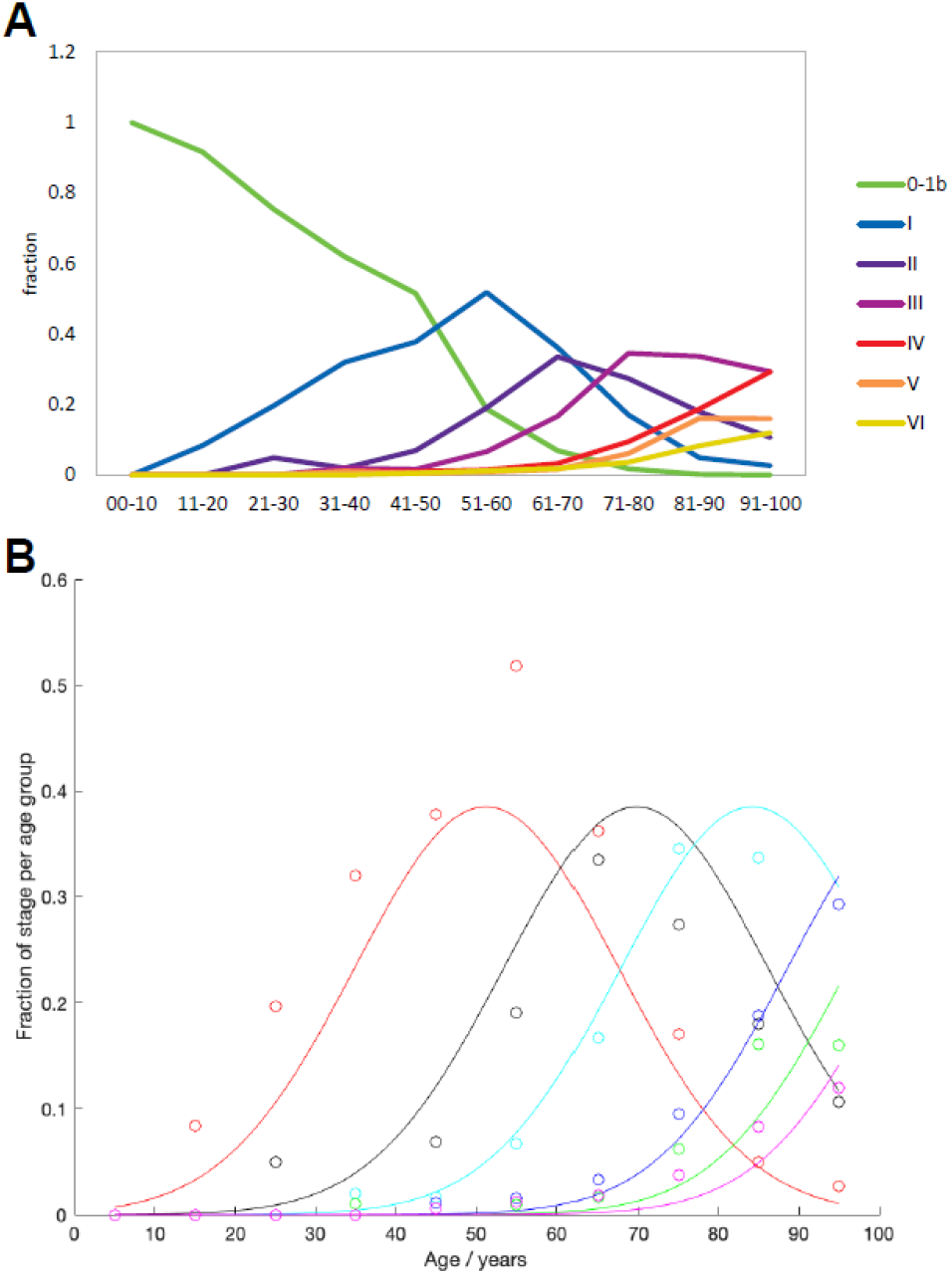
Fractions of individuals in a particular Braak stage for a given age. Data from Braak *et al.* (14). (A) All patients below Braak stage I have been grouped together into one group, the remainder are as classified by Braak *et al.* (B) The above distributions are fit to Gaussians, where the magnitude and standard deviations of the Gaussians are global parameters determined by the distributions of stages I - III and only the midpoint is a free parameter for all stages.

Simulations agree best with experiment if spreading is fast, Fig. 1e, notably also recovering the decrease in the EC and hippocampus, as observed previously (19). By contrast, if spreading is slow, the curves differ more significantly from those obtained in experiments, Fig. 1f. Crucially however, the fact that a significant concentration of seeds is present throughout widespread regions already at Braak stage III means that the system can never truly be spreading-limited; even if spreading is slow, the replication of seeds already present will dominate and no true propagating front emerges. Indeed, the simulations show that the time to reach a high seed concentrations in all regions shows little dependence on the spreading rate, but strongly depends on the replication rate (see SI section 2.2). To confirm the early presence of tau aggregates in the neocortical regions by an orthogonal method, we quantified aggregated tau in brain slices stained with the anti-tau antibody AT8 by image analysis. The data (Fig. 2b) confirm the presence of neuritic AT8 positive pathological staining can be detected even in the primary visual cortex at Braak stage III. Thus, the conclusion that local replication, not long-range spreading, is the process that dominates the overall kinetics of tau accumulation after Braak stage III is robust, and its inhibition would therefore have the most significant effect on delaying tau accumulation.

### Tau seeds in AD have a doubling time of approximately 5 years

Having thus established that the time it takes for tau to spread between brain regions does not influence the kinetics of tau seed accumulation, we can extract quantitative information from the measurements of seeds in AD brains in the form of an effective replication rate. We fit the approximate solution of equation (1) in the replication-limited case (see SI, equation 9) to the increase in seed concentration over time, measured by a range of methods, in a number of different neocortical regions, see Fig. 2.

For simplicity Fig. 2 shows the least square fits to the median, but Bayesian Inference on all datapoints was performed to determine the effective rate constant of replication as κ ≈ 0.17±0.05 years^−1^ (errors are one stan<u>dard</u> deviation, details see Methods).

This corresponds to a doubling time, *t*_2_, i.e. the time taken to double the number of seeds, of approximately 4 years, where 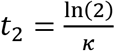. Our analysis of additional seeding data from Furman *et al.* (20) and Kaufman *et al.* (21), yields rates consistent with our data of *κ* ≈ 0.2±0.1 years^−1^ (t_2_ ≈ 3.5 years) and *κ* ≈ 0.08±0.02 years^−1^ (t_2_ ≈ 9 years), respectively (see Fig. 2c,d). Additionally, we also analyse data obtained by other measures of tau accumulation. (i) Staining of brain slices by an anti-tau antibody, AT8 (see Fig. 2b) followed by quantification by image automated analysis. (ii) Quantification of aggregated tau by ELISA, which was performed in addition to seed amplification measurements by Furman *et al.* (20). (iii) Stereological techniques to quantify the number of NFT in the banks of the superior temporal sulcus (high order association region) for individuals who had passed away a known number of years since developing symptoms, which was performed by Gomez-Isla *et al.* (22) who found a linear relationship between the number of accumulating tangles and the duration of disease. (vi) Longitudinal tau Positron Emission Tomography (PET) in human participants with varying levels of amyloidosis and clinical impairment, which was performed by Sanchez *et al*. (24) and evaluated longitudinal change in tau PET radio-tracer specific binding and measured change rates in individuals over the course of approximately 2 years.

While these measurements may quantify different forms of tau than the seeding assay, the same equations can be applied and we find replication rates of *κ* ≈ 0.17 years^−1^ (ELISA, t_2_ ≈ 4 years), *κ* ≈ 0.2 years^−1^ (AT8, t_2_ ≈ 3.5 years), *κ* ≈ 0.22 years^−1^ (stereological counting, t_2_ ≈ 3 years), *κ* ≈ 0.19 years^−1^ (PET, t_2_ 3:5 years). Crucially, this close agreement highlights that our findings are consistent across a variety of assays used to quantify aggregated tau. Combining the data (see Methods) yields an average rate of *κ* ≈ 0.14 years^−1^ corresponding to a doubling time of ~5 years. Moreover, while the time axis in the seed replication and ELISA data was obtained from the average time between Braak stages, the time in the stereological neuropathological counting data and the PET data corresponds to real time elapsed, since first occurrence of symptoms or time between PET measurements, respectively. This combination of a variety of methods to determine both the seed or aggregate concentration and the time confirms the robustness of our conclusions about the doubling time. Notably, all methods yield a remarkably low rate of replication, with a doubling time of several years, which implies that a large number of seeds and aggregates has to form initially in order to achieve the high final concentrations. Given the above rates, an increase by approximately 100-1000 fold is expected in the few decades of the disease, whereas the number of aggregates in the final stages is expected to be only a few orders of magnitude less than the number of neurons in the brain (i.e. tens of millions of aggregates, corresponding to one aggregate per every hundred neurons (22)), so would require an increase many orders of magnitude larger than that predicted if the disease were to be initiated by few aggregates in a specific location with a doubling time of 5 years. This implies that at Braak stage III there are many aggregates widely distributed across the brain.

To further compare these results with a common model system of tau pathology, we analysed data from P301S mice, measured by Holmes *et al.* (25) with the same seed amplification assay. Again, we find that spreading is not a rate-limiting step, i.e. aggregation either begins at many locations throughout the brain, as might be expected in a transgenic overexpressing animal, or begins at a small number of location but spreads much faster than the time-scale of the disease. Moreover, the increase in seeding activity is exponential during early disease, with a doubling time of approximately 2 weeks (details see SI section 1).

### Replication in AD is orders of magnitude slower than in mouse models or *in vitro*

The rates of replication for the systems analysed here are compared to the *in vitro* aggregation of tau, the prion protein and Aβ42 in Fig. 3b. To compute the relevant range of rates from the *in vitro* measurements of tau aggregation, we assumed that the monomer concentration was between 100 nM (estimate for monomeric tau in solution) and 10 μM (estimate for total monomeric tau) (26, 27). The differences between the individual systems are so large that they remain significant even given this uncertainty in the monomer concentration. We illustrate the biological meaning of the replication rates in the different systems by showing how long 36 rounds of doubling (producing ~70 billion seeds from one) would take in Fig. 3c. Measurements of the average size of tau seeds allow us to dissect the replication rate in humans derived above into the contributions from growth and from multiplication (for details see SI section 2.4) by using the fact that the replication rate κ is determined by the product of growth and multiplication, whereas the average size μ is determined by their ratio.

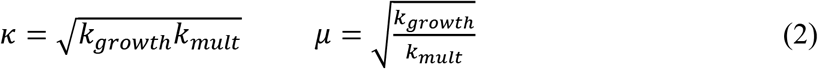

**Figure 3:**
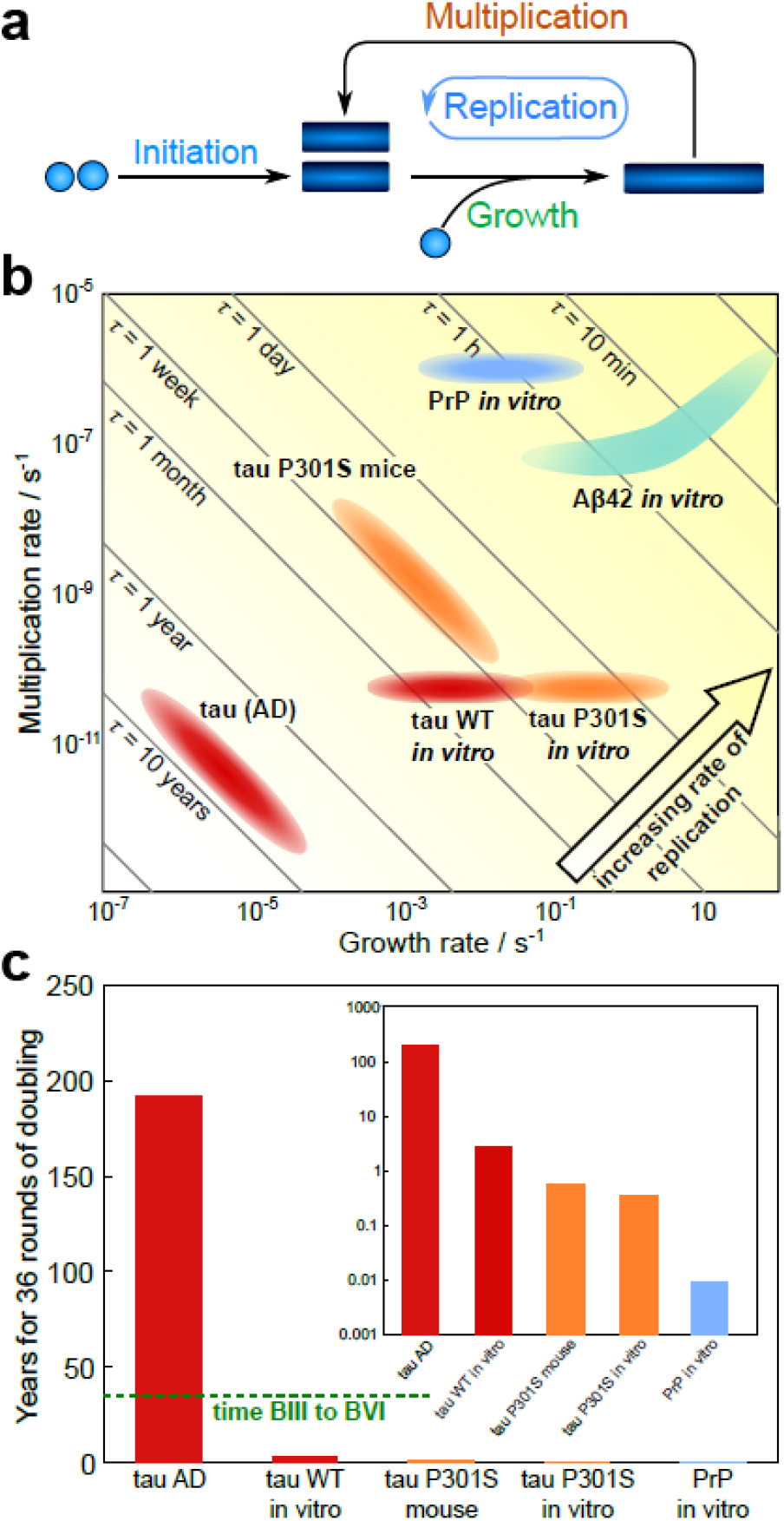
**a** Key steps of aggregation reactions, illustrating how growth and multiplication combine to lead to seed replication. **b** Comparison of the rates of tau seed growth and multiplication determined in AD brains here, with the rates predicted at concentrations between 100 nM and 10 μM from *in vitro* experiments (26), as well as the rates for both PrP (28) and A 42 (29) replication *in vitro* at the same concentrations (for details see Materials and Methods). Diagonal lines show the order of magnitude of the doubling time, i.e. points along the diagonal line have the same doubling time and replication rate. **c** Time required to produce 70 billion seeds form one seed (i.e. time for 36 rounds of doubling), for comparison with disease timescales, the dashed green line shows the average time that passes between Braak stage III and Braak stage VI. Only the two slowest systems, WT tau, are visible on a linear scale, inset shows time on a logarithmic axis.

The rates of both growth and multiplication are shown for the systems analysed here and compared to the *in vitro* aggregation of tau, the prion protein and A 42 in Fig. 3b. As can be seen in Fig. 3b, the replication rates of both PrP and Aβ42 are over an order of magnitude faster than those of tau *in vitro* or in mice, and tau replication in AD is even slower by a further two orders of magnitude than *in vitro*. Remarkably, while wild type tau replication in AD proceeds about two orders of magnitude slower than *in vitro*, the aggregation rate of P301S tau *in vitro* is comparable to replication in P301S mice. This is partly accounted for by the higher concentrations of tau in P301S mice, but nonetheless, the relative decrease of the rate *in vitro* compared to the rate measured *in vivo* is significantly less pronounced for P301S mice than for human AD. This observation may indicate that the mouse model lacks some of the mechanisms that inhibit tau aggregation in humans, especially in the setting of many fold over-expression of a protein that has high aggregation potential. Moreover, neurons may be more effective at preventing replication when it occurs more slowly, so that protective mechanisms are less easily overwhelmed.

The significantly lower rates of both growth and multiplication in humans highlight the importance of the mechanisms that have evolved in order to limit the rate of replication in living systems. The replication rate we determine here from human samples contains all these effects, from the clearance of seeds and aggregates by a variety of processes, to the effects of chaperones and other molecules that reduce the rates of aggregation and prevent self-replication of existing seeds. The relative slowing of growth and multiplication may serve as a guide to determine which mechanisms of inhibition are most prominent *in vivo*.

## Discussion

A key consideration in the interpretation of the results is the level of coarse-graining imposed by the experimental data, which affects the interpretation of the effective replication and spreading rates obtained from their analysis. The spatial resolution of the data is at the level of a single brain region, so while spreading between brain regions, referred to as long-range spreading above, can be recovered from these data, the effects of local inhomogeneities are subsumed into an effective replication term here. In other words, while it is clear that spreading between brain regions is not rate-limiting in the overall accumulation of tau seeds after Braak stage III, effects on lengthscales smaller than those resolved in the data, such as transfer between neighbouring cells within one region, may be of significance. Considerations of more spatially resolved data from model systems may serve as a guide as to which local transport processes, if any, are most likely to significantly contribute to the effective rate of replication that we determine from the coarse-grained data.

In particular, the current model does not explicitly distinguish slow transfer across a synapse / fast aggregation once it gets into the next cell from fast transfer and slow aggregation. In model systems that contain overexpressed mutant (but not wild type) tau, uptake and aggregation in the recipient cell can occur very quickly (within hours to days) (12,25). Yet injecting seed competent tau into mice that do not overexpress tau leads to production of AT8 positive phosphotau neuronal inclusions over many months (7), suggesting that the rapid aggregation seen in model systems may be a feature of high levels of very aggregation-prone substrate, which is unlikely to occur in the human condition. Therefore, consistent with the overall model we propose, local transport is unlikely to be slow enough to limit the rate and aggregate formation, at the synapse and in post-synaptic compartments, appears to be the slow step in NFT formation in human Alzheimer disease and thus our coarse-grained description of replication is indeed likely to represent the molecular process of seed replication.

The finding that replication is rate-limiting and that it proceeds quite slowly raises the question of how spatial inhomogeneities arise and how the high concentrations of seeds are created prior to Braak stage III. No definite conclusions on either question can be drawn given our current data, however, spatial and temporal variation in the rates, such as a differing propensity for initial seed formation, which is higher in the EC and surrounding regions, might explain the observed behaviours.

In conclusion, by applying chemical kinetics to *in vivo* data we were able to describe the spreading and replication of seed aggregates in the brain. Using data from human AD brains, we find that the process of tau seed accumulation is dominated by the local replication of seeds, and that spreading between brain regions appears not to be a rate-limiting step after Braak stage III, in the neocortical regions. The exponential increase in seed number observed in these regions is strong evidence that tau aggregates replicate auto-catalytically. From these data, we are able to extract the rate of tau seed replication in human AD and find that this rate is orders of magnitude slower than that measured for purified tau *in vitro*, quantifying the effectiveness of innate cellular mechanisms that curtail tau seed replication. Remarkably, we also find that the replication rate is so slow that the high numbers of seeds present in late disease require that either many seeds are formed de novo, rather than from existing seeds, or seed replication proceeds much faster prior to Braak stage III. The conclusions from our model, built from biosensor data and confirmed both by retrospective neuropathological analyses and prospective PET analyses, show that tau replication rather than transfer down an axon to the next neuron is likely to be the rate limiting step during the mid and later stages of AD, which has important implications for current therapeutic strategies. We envisage that the models developed here will form the basis for determining the rate-limiting processes and quantifying their rates for a wide range of other tauopathies, as well as aggregation-related neuro-degenerative diseases in general.

## Data availability statement

The data that support the findings of this study are available from the corresponding author upon reasonable request.

## Acknowledgments

We thank Michel Goedert and Will McEwan for useful discussions. We acknowledge funding from Sidney Sussex College Cambridge (GM) and the European Research Council Grant Number 669237 (to D.K.) and the Royal Society (to D.K.). The Cambridge Brain Bank is supported by the NIHR Cambridge Biomedical Research Centre.

## Author contributions

G.M and D.K. conceived the study; K.A. acquired the data; G.M., Y.Z., C.K.X. analysed the data; G.M., Y.Z., T.R., S.D., J.S., K.A.J., J.B.R., B.T.H., T.P.J.K., and D.K. interpreted the data; all authors contributed to the writing of the manuscript.

## Material and Methods

### Immunohistochemistry

Human brain tissue from 25 brain donors was obtained from the Cambridge Brain Bank (NRES 10/H0308/56). The donated brains had been pathologically assessed by a neuropathologist following the Consortium to Establish a Registry of Alzheimer’s Disease (CERAD) and Braak staging protocols. Cases were selected to include a variety of Braak stages. This included 7 Braak stage 0 (mean age 64 years; range 35-83), 8 Braak stage III (mean age 84 years; range 72-95) and 10 Braak stage VI (mean age 74 years; range 60-89).

Deparaffinized 10 μm sections of the primary visual cortex were obtained. These were subjected to antigen retrieval in 98% formic acid for 5 minutes followed by 4% aqueous hydrogen peroxide to block endogenous peroxidases. Sections were then rinsed with tap water and phosphate buffered saline (PBS) before being blocked with normal rabbit serum (Dako) in PBS.

Sections were then incubated with antibody to phosphorylated tau protein (AT8 1/500, Thermo) for 1 hour. After rinsing for 5 minutes in PBS they were incubated with secondary antibody (Rabbit Anti-Mouse 1/200, Dako) for 30 minutes. After rinsing for 5 minutes in PBS they were incubated in avidin-biotin complex (ABC, Vector) for 30 minutes before being developed with diaminobenzidine (DAB, Vector). Slides were then lightly counterstained with haematoxylin.

Digital images were obtained using a camera (Infinity 2, Lumenera) attached to a micro-scope (Olympus BX53). Images of the primary visual cortex were obtained at x200 magnifica-tion to create images measuring 5.892 mm^2^.

### Linking Braak stage to time since disease onset

While in mouse models the time since disease onset can simply determined from the age of the animal, in human patients this becomes more difficult. The age of onset varies over decades and symptoms only appear in the later stages, meaning that samples of early disease stages are from patients that often had not been diagnosed prior to their death. The disease stage is determined by inspection of post-mortem brains, which allows classification into different Braak stages.

Here we attempt to link Braak stage to time since disease onset to then be able to put the measurements of seeding activity on a common time axis. We use data on the age and Braak stage of 2332 individuals, published by Braak *et al.* (14). The underlying assumption required to proceed is that once the disease has begun, it progresses in much the same manner in different individuals, i.e. the differences between individuals originate mainly from the different times of disease onset, the disease progression itself is less variable. If this was indeed the case, one would expect the age-distribution for each disease stage to have approximately the same shape, but a different average age. Moreover, the mean or median age at each stage should be a good guide to determine the average time spent in each stage.

The data by Braak *et al.* (14) have 10 age categories, each spanning a decade, between 1 and 100 years, and 12 Braak sub-stage, from stage 0 to stage VI. We initially normalise the data for each age group, resulting in the probability of being in a certain Braak stage, given the age.

During early Braak stages (1a to III) the distributions look Gaussian and can easily be fitted. From stage III onwards, the fact that there are no data available for people above 100 years of age means that the distributions are cut off. To deal with this problem, we fix the magnitude and standard deviations of the Gaussians based on the early stages and only fit the mean for the later stages of disease. The means, i.e. predicted average ages for each stage are shown in table 1. An alternative measure for the timescale of disease is provided by Whittington *et al.* (30) who obtained a measure of amyloid load as a function of time during AD using PET imaging. Their results show a progressive increase of amyloid load over approximately 30 years, consistent with the timescales we here obtain for transitioning from Braak stage III to Braak stage VI.

**Table 1:** Predicted mean age for each Braak stage and difference between consecutive stages.

| Stage | Mean age | Time to next stage |
| --- | --- | --- |
| 0-Ib | 19 | 32 |
| I | 51 | 19 |
| II | 70 | 14 |
| III | 84 | 21 |
| IV | 105 | 8 |
| V | 113 | 5 |
| VI | 118 | - |

### Fitting to obtain replication rates

To obtain the effective replication rate, we use a rescaled version of equation 9, describing the measured values:

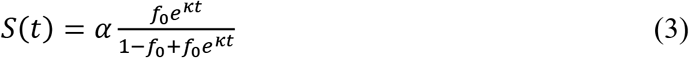

where *α* is the parameter that converts fraction of seeds, *f(t)*, to the measured quantity (e.g. intensity of AT8 response, ELISA signal etc.). For clarity, we show least squares fits to the averages for the data points figure 2 but we use Bayesian inference on the individual measurements, assuming normally distributed noise, with the standard deviation fixed from the standard deviation of repeat measurements. We assumed a flat prior for *α* and a *1/x* prior for *κ* and *f_0_*. The bounds for the initial fraction of seeds, *f_0_*, were chosen to be between 10^−10^ and 0.01, corresponding to there being on the order of one seed per brain and to there being 1% of the final seed concentration at the beginning of the disease, respectively. We believe these are very generous bounds and any values outside them are very unlikely. Where available, the measurements for each brain region were analysed separately, the resulting posterior was marginalised over *α* and *f_0_*, yielding a posterior distribution for *κ*. For datasets with more than one brain region, these marginalised posteriors were then combined to give an overall posterior for *κ*, for the entire dataset. We furthermore combined the posteriors of all experiments (different datasets of seed measurements, AT8 quantification, ELISA and stereoscopic counting) to yield an over-all value for *κ*, which we use to represent tau in AD in Fig. 3. This value can be interpreted as the value of *κ* most consistent with all data recorded, by all different methods.

### PET data and analysis

Unlike the other datasets analysed, which had both a measure of the time since disease onset and a measure of the the seed concentration, the PET data from Sanchez *et al.* (24) instead provide a measure of the seed concentration at two timepoints in the same patient, separated by approximately 2 years. The analysis of these data was performed by assuming the tau PET signal follows

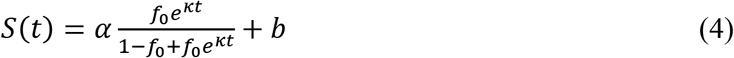

where the parameters are as defined in equation 3 and the additional parameter *b* accounts for the non-zero baseline. The two consecutive measurements allow the determination of an annual rate of increase *r*, which we take as an estimate for the time derivative, *r* ≈ Ṡ(*t*).

Differentiating and rearranging equation 4 yields the replication rate *κ* in terms of the PET signal, *S(t)*, and the rate, *r*

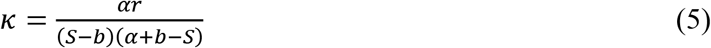

where *S* is the PET signal averaged over the consecutive measurements. Fig. 2 shows a plot of *r* against *S*. From the median values of patients with mild cognitive impairment (MCI) and with AD, we obtain *κ* = 0.19 years^−1^ and thus a doubling time of approximately 4 years. We chose *b* = 1.19, based on the PET signal in healthy patients and *α* = 2.31 based on the maximal signal observed as well as the curve-shape in Fig. 2. The solid line in Fig. 2 is the predicted behaviour based on this value of *κ*. The data for cognitively normal patients with a high Aβ load (as determined by a high signal in ^11^C-Pittsburgh Compound B PET measurements, details see Sanchez *et al.* (24)) are also shown but not included in this analysis. They produce *κ* = 0.37.

### Comparison of *in vivo* and *in vitro* rates

*In vitro*, we have control over the concentrations of the reaction species and are therefore usually able to determine rate constants, rather than just rates, which in turn allows us to extrapolate to predict the rates at different concentrations. Given the rate constants from previously published work, we here evaluate the rates at a range of monomer concentrations to obtain a range of relevant values for comparison with the rates measured *in vivo*. We estimate the relevant range of tau concentrations in AD to be 100 nM to 10 μM, and thus we evaluate the rates based on *in vitro* measurements in this range of concentrations. We also evaluate the rates of PrP and A aggregation in this range, simply to provide a reference on relative speed compared to other proteins. The *in vitro* experiments, in which the rate constants used here were determined (26, 28, 29), were performed at μM concentrations and are thus in the relevant range, making errors from extrapolation small. The region predicted for A 42 in Fig. 3 is curved because its multiplication rate depends on concentration.

### Estimates of effective diffusion constants

In SI section 2.1 we show that for systems with a compact initial distribution the switch between the replication-limited and the spreading-limited regimes occurs approximately when *D/κ* = 0.0025 in reduced units obtained by setting *r*_max_ = 1. To obtain an approximate value for the effective diffusion constant in the context of tau seeds in AD, we set *r*_max_ = 10 cm and *κ* = 0.15 years^−1^, where we estimated the distance that seeds have to travel to reach any brain region from the region of initial seed formation to be 10 cm and the replication rate *κ* was obtained from fits of experimental data. The effective diffusion constant at which the switch from replication limited to spreading limited occurs is then approximately *D* ≈ 10^−13^ m^2^s^−1^. Diffusion coefficients for proteins are generally on the order of 10^11^ m^2^s^−1^. Therefore, considering the fact that the effective diffusion constant we calculate here is for seeds, which are likely significantly larger than individual proteins, and that it may include both active transport processes and membrane crossing, values either side of this critical value of *D* ≈ 10^−13^ m^2^s^−1^ are entirely reasonable. In the simulations of data from AD brains, Fig. 1e,f, effective diffusion constants of *D* = 0.025 (fast spread) and *D* = 0.00025 (slow spread), with a replication rate *κ* = 0.15 years^−1^ were used. In analogy to the above calculation, these would correspond to *D* ≈ 10^−11^ m^2^s^−1^ and *D* ≈ 10^−13^ m^2^s^−1^ respectively. Note that these values of *D* are slightly different from the above values because the initial distributions used in the simulations starting from Braak stage III can no longer be considered a compact distribution near the origin. Alternatively, to avoid the assumption of a specific length-scale, we can express the length in the same way as in the experimental data, in terms of number of crossings between brain regions, assuming there are 6 regions corresponding to each Braak stage. In that case the fast and slow spread corresponds to *D* ≈ 1 (regions)^2^year^−1^ and *D* ≈ 0.01 (regions)^2^year^−1^. In other words, the average time for a seed to cross from one region to the next is 1 and 100 years in the fast and slow spread limits. Therefore, even when the average time for a given seed to cross from one region to the next is a year, spreading would be considered fast compared to replication.

