## Supplementary Information for "Amplification, not spreading limits rate of tau aggregate accumulation in Alzheimer’s disease"

#### 1 Additional data

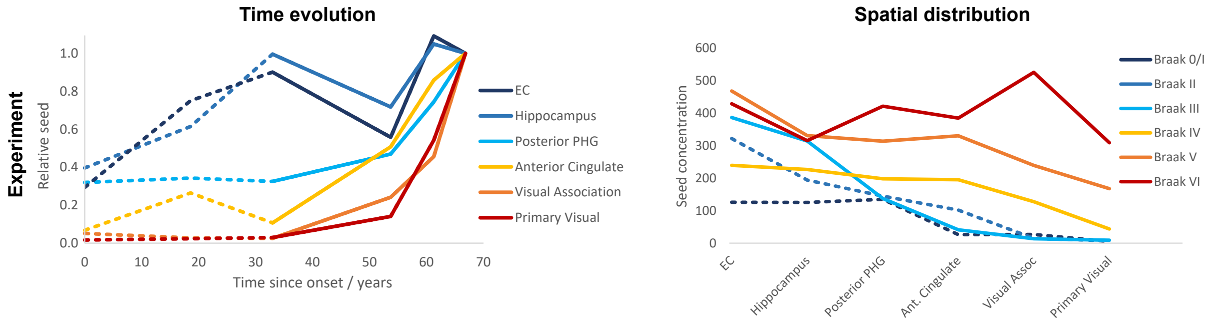


Figure S1: Full data from DeVos et al.[16], the main text shows only Braak stages 3 onwards (solid lines).

**Sensitive tau seed measurements in mouse models show that behaviour is limited by replication, not spread.** Utilising the ability of tau seeds to replicate, Holmes et al.[2] developed a sensitive assay for the determination of tau seeds by FRET Flow Cytometry and measured the seeding activity of tau aggregates in several brain regions of P301S transgenic mice, at several times up to one year of age. The data were extracted from Holmes et al.[2], Figures 1c and 5 in their work. The data in the former was used to obtain a calibration curve, which was then used to convert the data from the latter from FRET intensity to seed concentration. The data and fitted calibration curve are shown in Fig. S2. Several of the data points in Holmes et al. figure 5 lie outside this calibration range, and were thus excluded from Fig. S3.

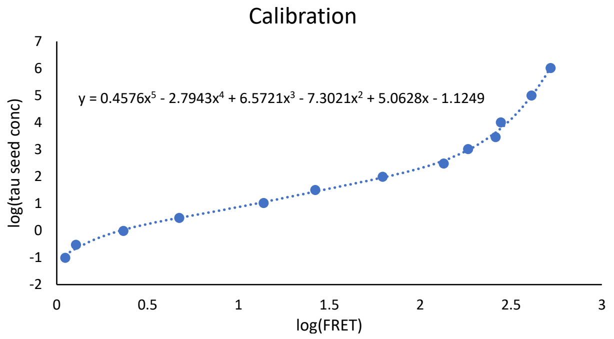


Figure S2: Data from Figure 1c in Holmes et al.[2] (dots), used to obtain a calibration curve (dashed line) that allows conversion of FRET intensity to tau seed concentration. The fifth order polynomial used is given in the figure.

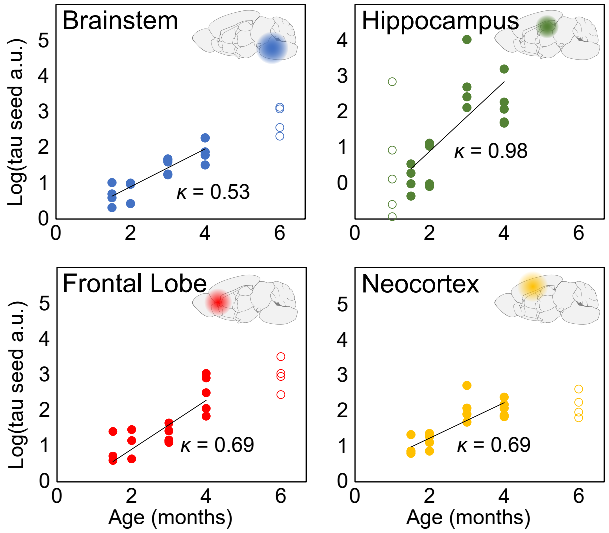


Figure S3: The logarithm of tau seed concentration in P301S transgenic mice, measured by FRET Flow Cytometry, at several time-points and locations throughout the brain. The data (filled and open circles) were obtained from Holmes *et al.* [2]. The solid line is a straight line fit in logarithmic space. For the fitting, only times were taken into account for which data exist in all brain regions (filled circles) to avoid biases.

Two features of these data are noteworthy: First, there is no evidence of a delayed initiation of the aggregation reaction in any brain regions, Fig. S3. Therefore, spreading is not a rate-limiting step. Second, the increase in seeding activity is exponential during early disease. This is indicative of an exponential increase in the mass of aggregates, a hallmark of aggregate multiplication[31] and further evidence that multiplication of aggregates is also occurring *in vivo*. Analysing the data up to 4 months, we determine the rate of replication, $\kappa = 0.6$ months$^{-1}$, which corresponds to a doubling time of approximately 2 weeks (in the brainstem, neocortex and frontal lobe). In summary, the data acquired by Holmes *et al.* [2] show that their mouse models are clearly replication-limited and the spreading of aggregates from one brain region to the next is not a rate-determining step. Broadly speaking, two scenarios could give rise to a replication-limited behaviour. (1) The initial seeds are formed in one brain region only and spreading is so fast that it does not affect the kinetics, or (2) seeds are formed throughout the brain, so no long-range spreading is required. Thus, this kind of spreading may occur and even constitute a necessary step in disease progression, however, it does not limit the kinetics of tau seed accumulation.

**Rates of diffusional uptake from cell models** Evans et al. [32] detect the efficient uptake of aggregated tau (at nM concentrations) into human cortical neurons in cell culture over the timescale of hours. Similarly McEwan et al. [33] find that HEK293 cells expressing aggregation-prone tau take up seeds that trigger intra-cellular aggregation within hours, but also show that there are effective mechanisms to abolish the seeding effect once an aggregate has entered the cell. Together these results show that seed-competent aggregates can easily enter cells, but that there may be mechanisms that slow their replication, in agreement with our findings in this work.

### 2 Theoretical Models

#### 2.1 Reaction diffusion equation and the spreading-limited and replication-limited regimes

The equation describing spatially-dependent aggregation is:

$$\frac{\partial P(\mathbf{r},t)}{\partial t} = D\nabla^2 P(\mathbf{r},t) + kP(\mathbf{r},t)(P_{\max} - P(\mathbf{r},t)) \tag{S1}$$

where $P(\mathbf{r},t)$ is the aggregate concentration at time $t$ and position $\mathbf{r}$. $D$ is an effective diffusion coefficient and $k$ is an effective replication rate constant. The equation for the fractional seed concentration, $f(\mathbf{r},t)$, relative to the maximal seed concentration, eq. 1, is obtained by dividing the above by $P_{\max}$ and replacing $k = \kappa/P_{\max}$ and $P(\mathbf{r},t) = f(\mathbf{r},t)P_{\max}$. Fisher equation (i.e. the 1-dimensional version of eq. 1), for compact initial conditions such as a finite seed concentration at the origin, exhibits travelling wave solutions whose wave speed approaches $v = 2\sqrt{\kappa D}$.

We have not yet specified the dimensionality of the problem. Both a 3-dimensional model and a 1-dimensional model are reasonable: A 1-dimensional can be used to model axonal spreading, whereas a 3-dimensional one describes uniform spreading in space. More generally, in this problem the dimensionality simply encodes how more space is available as seeds spread away from the origin. Axonal spread on a branching network may best be described with a dimensionality between 1 and 3. Therefore, we here consider both extreme cases, 1-dimensional spreading and 3-dimensional spreading, and show that the qualitative results are the same. More importantly, in the replication-limited regime both models yield the same answer. As the accumulation of tau seeds in AD occurs in the replication-limited regime, the rate constants we obtain from experimental measurements are thus independent of the dimensionality of the model chosen here.

In 3 dimensions we consider the spherically symmetric problem (so $\nabla^2 = \frac{1}{r^2}\frac{\partial^2}{\partial r^2}r^2$ , where $r$ is the distance from the origin) with an initial concentration of aggregates at the origin and the boundary condition of a solid wall at distance $r = r_{\max}$ (i.e. no flux through the surface at $r = r_{\max}$). The only difference to the 1 dimensional system is thus that $\nabla^2 = \frac{\partial^2}{\partial r^2}$. Without loss of generality we set $r_{\max} = 1$ thus defining distances in units of the maximum distance from the origin $r_{\max}$. In 3 dimensions

$$\frac{\partial f(r,t)}{\partial t} = D\frac{1}{r^2}\frac{\partial}{\partial r^2}r^2 f(r,t) + \kappa f(r,t)\left[1 - f(r,t)\right] \tag{S2}$$

and in 1 dimension

$$\frac{\partial f(r,t)}{\partial t} = D\frac{\partial}{\partial r^2}r^2 f(r,t) + \kappa f(r,t)\left[1 - f(r,t)\right] \tag{S3}$$

**Limiting analytical solutions.** In the replication limit, the behaviour becomes independent of location, so an approximate solution to equation S2 can be obtained by setting the derivatives with respect to the spatial components to zero, yielding

$$f(\mathbf{r},t) = \frac{\langle f_0\rangle e^{\kappa t}}{1 - \langle f_0\rangle + \langle f_0\rangle e^{\kappa t}} \tag{S4}$$

where $\langle f_0\rangle$ is the initial concentration averaged over all space. This approximation is the same regardless of dimension.

In the spreading limit, the initial profile of increase is propagated throughout the reaction volume with speed $v$. For the 1-dimensional Fisher equation, it has been shown that $v = 2\sqrt{D\kappa}$ for sufficiently sharp initial distributions. We will use this speed here to obtain an approximate solution in both the 1-dimensional and the 3-dimensional case. The approximate solution is

$$f(r,t) = \frac{f_0 e^{\kappa(t+r/v)}}{1 - f_0 + f_0 e^{\kappa(t+r/v)}} \tag{S5}$$

where $f_0$ is the initial fraction of seeds at the origin. This approximation does not include the time delay due to diffusion of seeds from the initial distribution to establish the shape of the propagating front, so it performs less well for sharp initial distributions. This solution does not feature in the data analysis of this work, we merely include it here for completion. More accurate solutions can be found in the extensive literature studying the Fisher equation. A comparison of these approximate solutions and the numerical integration of equation S2 is shown in Fig. S4.

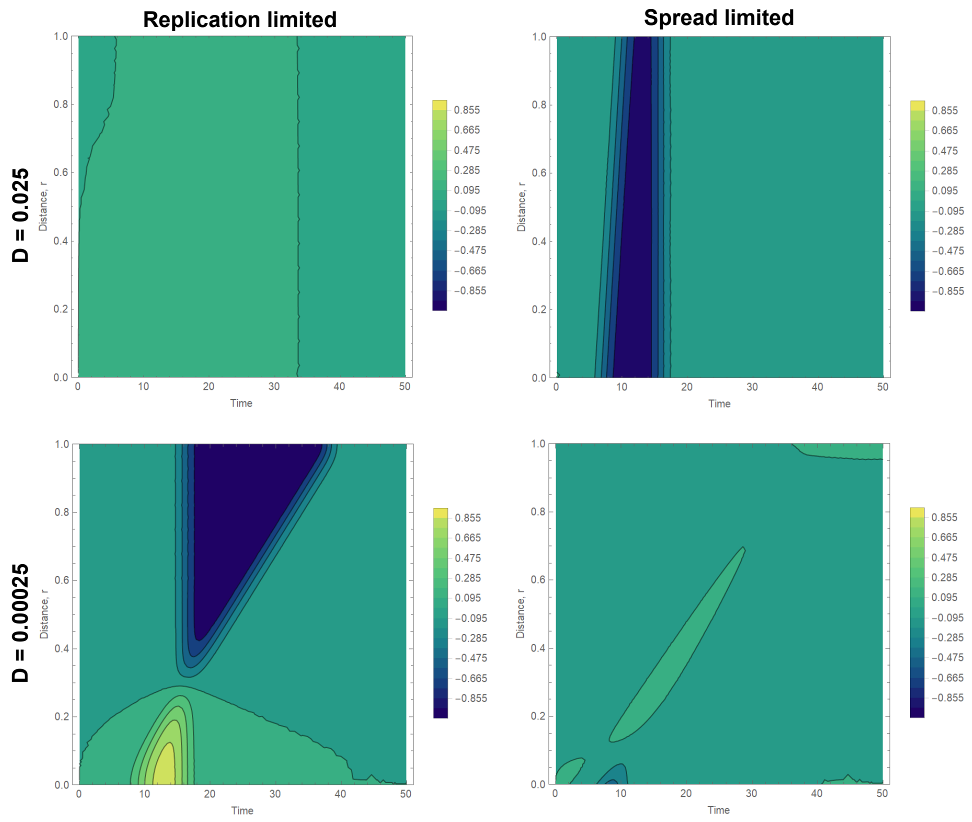


Figure S4: Difference between the numerical solution to equation S2 and the approximations (numerical - approximate), against distance and time. Green corresponds to good agreement, blue to an overestimation of the numerical solution and yellow to an underestimation of the numerical solution. The initial distribution was chosen to be $P(r, 0) = 0.1$ for $r < 0.01$ and 0 elsewhere. The approximation obtained in the replication limit, equation S4, performs well for fast diffusion $D \gg 0.0025$, top row. The approximation obtained in the spreading limit, equation S5, performs better for slow diffusion $D \ll 0.0025$, reproducing the correct spreading velocity. However, the time taken to initially establish the travelling wave leads to a delay and slight offset of the numerical solution to our approximation.

**Transition between regimes.** A transition between the spreading-limited and replication-limited regimes occurs when the time taken to spread throughout the entire reaction volume is comparable to the time needed to reach the maximal aggregate concentration at a given point, i.e. when $r_{\text{max}}/v \approx \tau_r$, where $\tau_r$ is the time-scale of the local reaction. We again approximate $v$ by the speed of a travelling wave solution to the Fisher equation, $v \approx 2\sqrt{\kappa D}$, thus we obtain in reduced units

$$\kappa D_{\text{switch}} \approx \frac{1}{4\tau_r^2} \tag{S6}$$

What remains is to estimate the time-scale of the local reaction $\tau_r$. Using equation S4 to approximate the local behaviour, we obtain

$$\kappa\tau_r = \log\left(\frac{(1-f_{\text{init}})}{f_{\text{init}}}\right) + \log\left(\frac{f_{\text{end}}}{(1-f_{\text{end}})}\right) \tag{S7}$$

where $f_{\text{init}}$ and $f_{\text{end}}$ are the local concentrations at the beginning of the reaction and when it is considered completed, respectively. To estimate $\tau_r$, these parameters need to be chosen. Their choice is to some extent arbitrary although for reasonable values the exact choice has little effect on $\tau_r$. We choose $P_{\text{end}} = 0.99$ and $P_{\text{init}} = 0.001$, giving a replication time-scale on the order of $\kappa\tau_r \approx 10$. Thus at the switch we have $\frac{D_{\text{switch}}}{\kappa} \approx 0.0025$. Indeed, numerical integration of equation S2, setting $\kappa = 1$ to set units of time, does show the expected behaviour in the two limits and a switch at $D \approx 0.0025$, see main text Fig. 1.

**Time evolution of total aggregate mass and importance of spatially resolved data.** Given the numerical solution to equation S2 for all positions and times, one can moreover calculate the time evolution of total aggregate mass, a measure that may be of importance if the available data are not resolved spatially. While we do not use these results in the main text, we present them here for completeness and to illustrate the importance of spatially resolved data in determining the mechanism. The total aggregate mass is given by an integration of the aggregate concentration over all space. In 3 dimensions the problem is spherically symmetric so we get

$$M_{\text{tot}} = \int_0^1 4\pi r^2 f(t,r)dr \tag{S8}$$

which can be evaluated from the numerical solution (in 1 dimension the factor of $4\pi r^2$ is missing). In the spreading and replication limits however, we can obtain approximate solutions. In the replication limit, the system is spatially uniform, thus the total aggregate mass follows the same functional form as the local increase, regardless of dimensionality,

$$M_{\text{tot}} = P_{\max} V \frac{e^{\kappa t} f_0}{1 - f_0 + e^{\kappa t} f_0} \tag{S9}$$

with $f_0$ being calculated from the average initial concentration, assuming it has diffused through the entire reaction volume. The pre-factor $V$ simply accounts for the total reaction volume with $V = \frac{4\pi}{3}$ in 3 dimensions and $V = 1$ in one dimension.

In the spreading limit, we assume that the concentration is at its maximum inside the propagating front, and is 0 outside. The radius of region inside the propagating front at time $t$ is given by $r = v * t$, where again we approximate the velocity of the travelling wave by the result obtained from Fisher's equation for compact initial distributions, $v = 2\sqrt{D\kappa}$. Thus in 3 dimensions

$$M_{\text{tot}} = P_{\max}\frac{4\pi r^3}{3} = P_{\max}\frac{32\pi}{3}(D\kappa)^{3/2}t^3 \tag{S10}$$

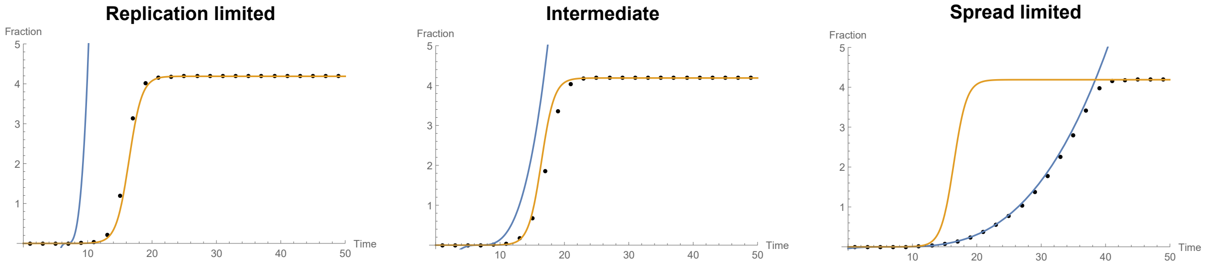


Figure S5: Time evolution of the total aggregate mass in the replication limited regime ($D = 0.025$), an intermediate regime ($D = 0.0025$) and the spreading limited regime ($D = 0.00025$), for an intitial distribution $P(r, t = 0) = 0.1$ for $r < 0.01$ and 0 otherwise. The black dots are derived from numerical integration of equation S2, the orange line is the approximation derived in the replication limit, equation S9, the blue line is the approximation derived in the spreading limit, equation S11.

until time $t = 1/v$, when the front reaches the boundary and the aggregate mass plateaus at $M_{\text{tot}} = \frac{4\pi}{3}$ (in 1 dimension this trivially becomes $M_{\text{tot}}/P_{\text{max}} = r = 2\sqrt{D\kappa t}$ for $t < 1/v$ and 1 otherwise). However, before the front starts propagating, it may have to establish the shape of the propagating front and may have to locally replicate to the maximum concentration. To approximate this we estimate the waiting time to get to 99% of the maximal aggregate concentration $t_c = \log(\frac{0.99(1-u_0)}{0.01u_0})$ where now $u_0$ is the local initial concentration. So overall we get

$$M_{\text{tot}} = P_{\text{max}} \frac{4\pi r^3}{3} = P_{\text{max}} \frac{32\pi}{3} D^{3/2}(t - t_c)^3 \qquad \text{(S11)}$$

and $M_{\text{tot}} = 0$ for $t < t_c$ (and analogous for 1 dimension). These solutions are compared with the numerical results for 3 dimensions in Fig. S5 and found to agree well. The increase of total aggregate mass is more sudden in a replication-limited system, however, in particular if the initial concentration is low and thus $t_c$ is large, the curves produced from spreading-limited behaviour in the 3 dimensional model can appear similar in shape. Thus, depending on the level of noise, it may be difficult to distinguish between the two cases given measurements of the total concentration only, and spatially resolved experiments may often be required to verify which process limits the overall rate.

#### 2.2 Effect of initial distribution in each limit

Here we illustrate the effect of lowering either the effective diffusion constant $D$ or the replication rate $\kappa$ by a factor of 3. While we label the limiting behaviours as spread-limited and replication-limited, even in the spreading-limited regime, the velocity of the propagating front depends on the replication rate via $v \approx 2\sqrt{\kappa D}$, thus a decrease of $\kappa$ always decreases the overall rate. We compare the case where the initial distribution is sharp and confined to a region close to the origin ($P(r, 0) = 1$ for $r < 0.01$ and 0 otherwise) to the case where the initial distribution has most of its weight around the origin but has significant seed concentrations everywhere (modelling the distribution of seeds measured at Braak stage III).

In the replication-limited regime, Figs. S6 and S7, reducing $\kappa$ significantly reduces the rate at which new aggregates accumulate, regardless of initial distribution and dimensionality. Reducing $D$ has little effect, except for a sharp initial distribution, where this decrease pushes the system more towards a spreading limited regime (if the simulations are performed at higher values of $D$ as a reference, this effect disappears). The choice of dimensionality has a clear effect in this regime, mainly because the dimensionality determines what average concentration the initial distribution corresponds to. For example, the sharp initial distribution ($P(r, 0) = 1$ for $r < 0.01$ and 0 otherwise) gives an average of $\langle P_0 \rangle = 1 * 0.1/1 = 0.1$ in 1 dimension but an average of $\langle P_0 \rangle = 1 * 0.1^3/1 = 0.001$ in 3 dimensions. In the spreading-limited regime, Figs. S8 and S9, reducing $\kappa$ decreases both the steepness and the speed of the propagating front and reducing $D$ decreases the speed but increases the steepness for a sharp initial distribution. The 1-dimensional and the 3-dimensional systems behave very similarly. The effect of a decrease of either parameter is comparable, as expected. However, for the more spread out initial distribution, a decrease of $\kappa$ significantly delays the reaction while a decrease of $D$ has no apparent effect. This somewhat counter-intuitive observation originates in the fact that there are seeds present throughout the reaction volume already at early times: once the spreading rate decreases beyond a certain limit, the overall behaviour is again governed by the local replication of seeds. In other words, at very high spreading rates the system becomes independent of the spreading because seeds are quickly distributed throughout the volume and the rate of this distribution does not limit the overall rate. However, given the spread out initial distribution, the overall rate also becomes independent of the spreading rate at very low rates of spreading. because then the seeds already present locally replicate much quicker than new seeds can move into the region by spreading. Spreading is not required to allow this reaction to go to completion. The latter situation cannot arise for sharp distributions, where spreading is absolutely necessary because there are regions without any seeds, which can never reach completion without obtaining seeds by spread from other regions. In summary, while the overall rate in systems with a sharp initial distribution will depend on the spreading once the spreading rate is below a certain limit, the overall rate in systems with a more spread-out initial distribution is independent of the spreading both at low and high rates of spreading.

The key conclusion from this analysis are the following

- In all cases, even for a sharp initial distribution that leads to a clear spreading-limited behaviour, reducing replication is the most effective strategy to prevent accumulation of aggregates.
- Spread-out distributions are never in a truly spreading limited regime.

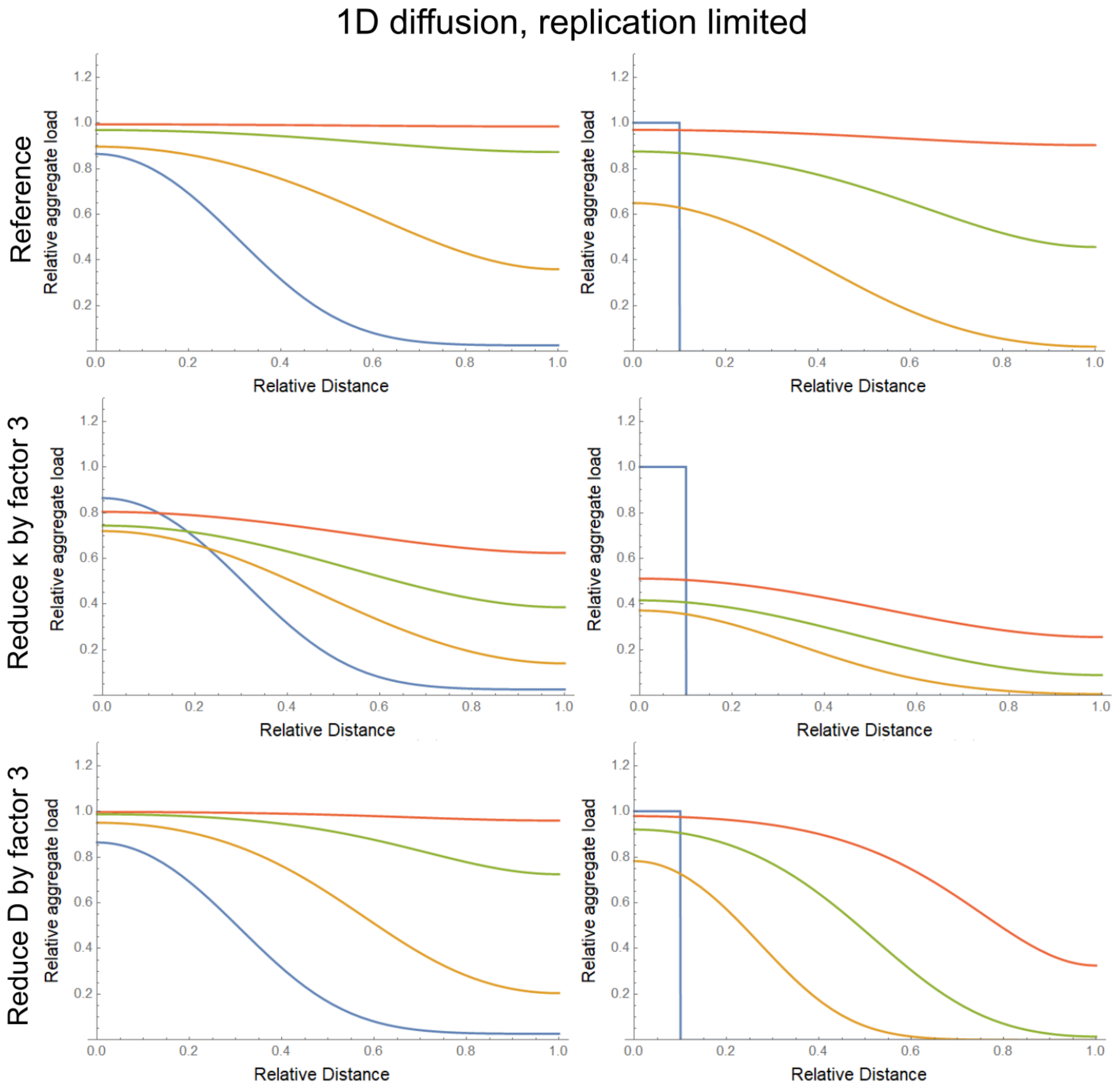


Figure S6: Effect of reduction in $D$ or $\kappa$ in the replication limited regime for 1-dimensional spreading, from numerical integration of equation S3. Reference values are $D = 0.025$, $\kappa = 1$ in reduced units. Times (0, 2, 4 and 6 in reduced units) are chosen to cover the range of behaviours.

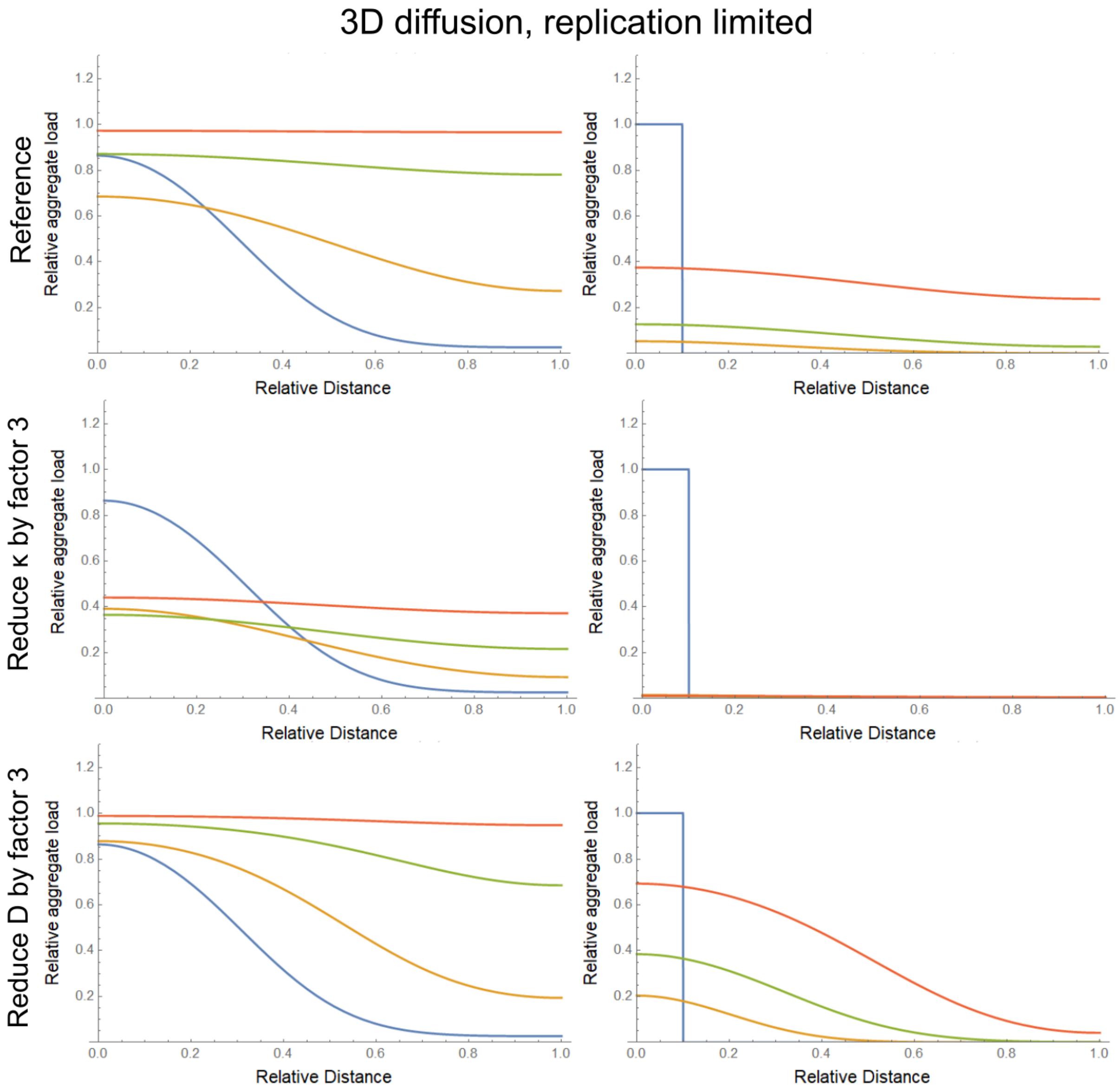


Figure S7: Effect of reduction in $D$ or $\kappa$ in the replication limited regime for 3-dimensional spreading, from numerical integration of equation S2. Reference values are $D = 0.025$, $\kappa = 1$ in reduced units. Times (0, 2, 4 and 6 in reduced units) are chosen to cover the range of behaviours.

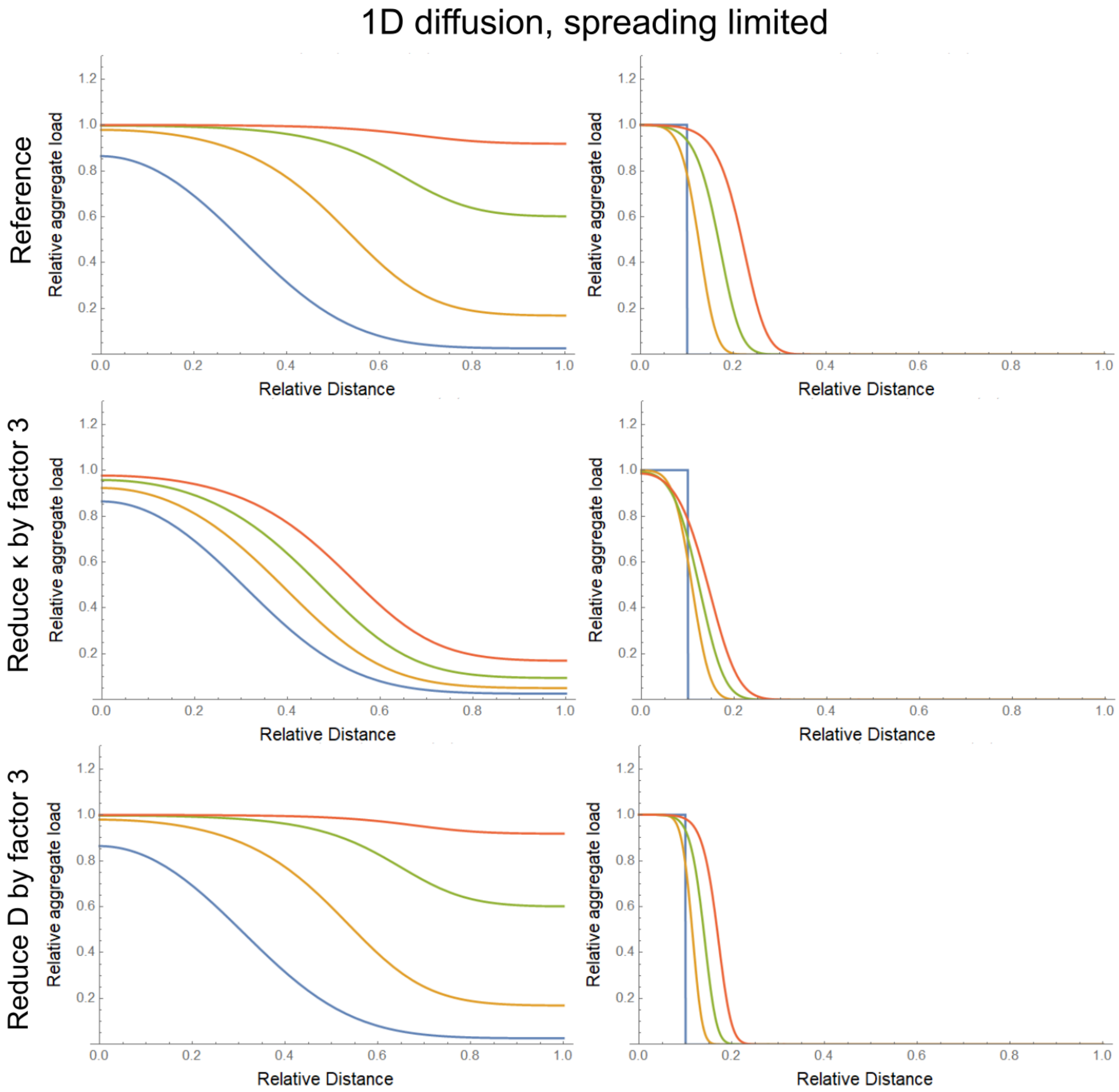


Figure S8: Effect of reduction in $D$ or $\kappa$ in the spreading limited regime for 1-dimensional spreading, from numerical integration of equation S3. Reference values are $D = 0.00025$, $\kappa = 1$ in reduced units. Times (0, 2, 4 and 6 in reduced units) are chosen to cover the range of behaviours.

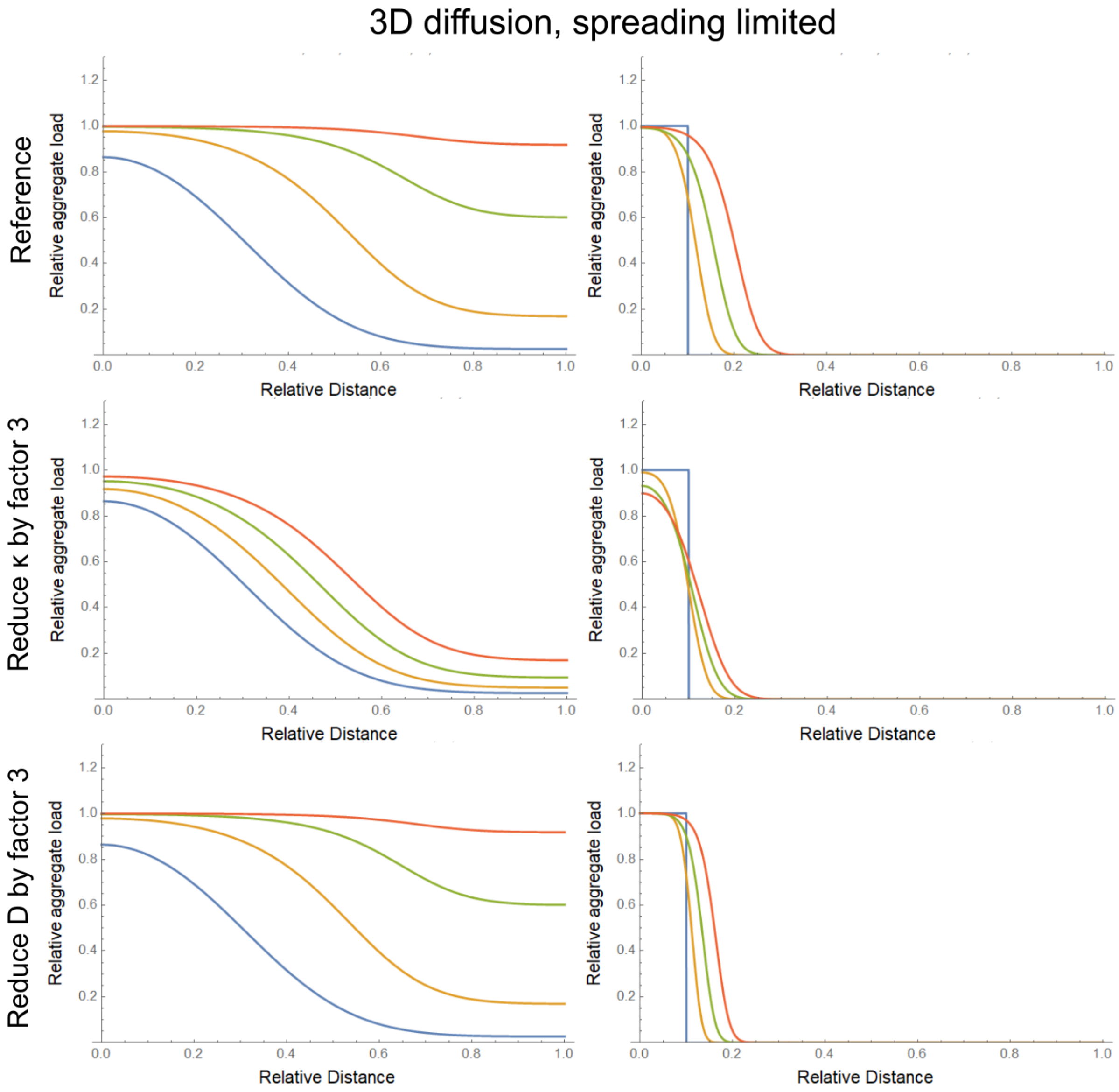


Figure S9: Effect of reduction in $D$ or $\kappa$ in the spreading limited regime for 3-dimensional spreading, from numerical integration of equation S2. Reference values are $D = 0.00025$, $\kappa = 1$ in reduced units. Times (0, 2, 4 and 6 in reduced units) are chosen to cover the range of behaviours.

#### 2.3 Effect of dimensionality - Spread through space and along axons

The simulations shown in Fig.1e,f are repeated in fig. S10 for both the 3-dimensional and the 1-dimensional model. The data are matched to a comparable degree in both cases, thus our conclusions hold regardless of the dimensionality of the model assumed. In practice this means that regardless of how much more volume is available to seeds the further away from the initially affected regions they are, we can conclude that the overall rate is limited by local replication, rather than spreading between brain regions.

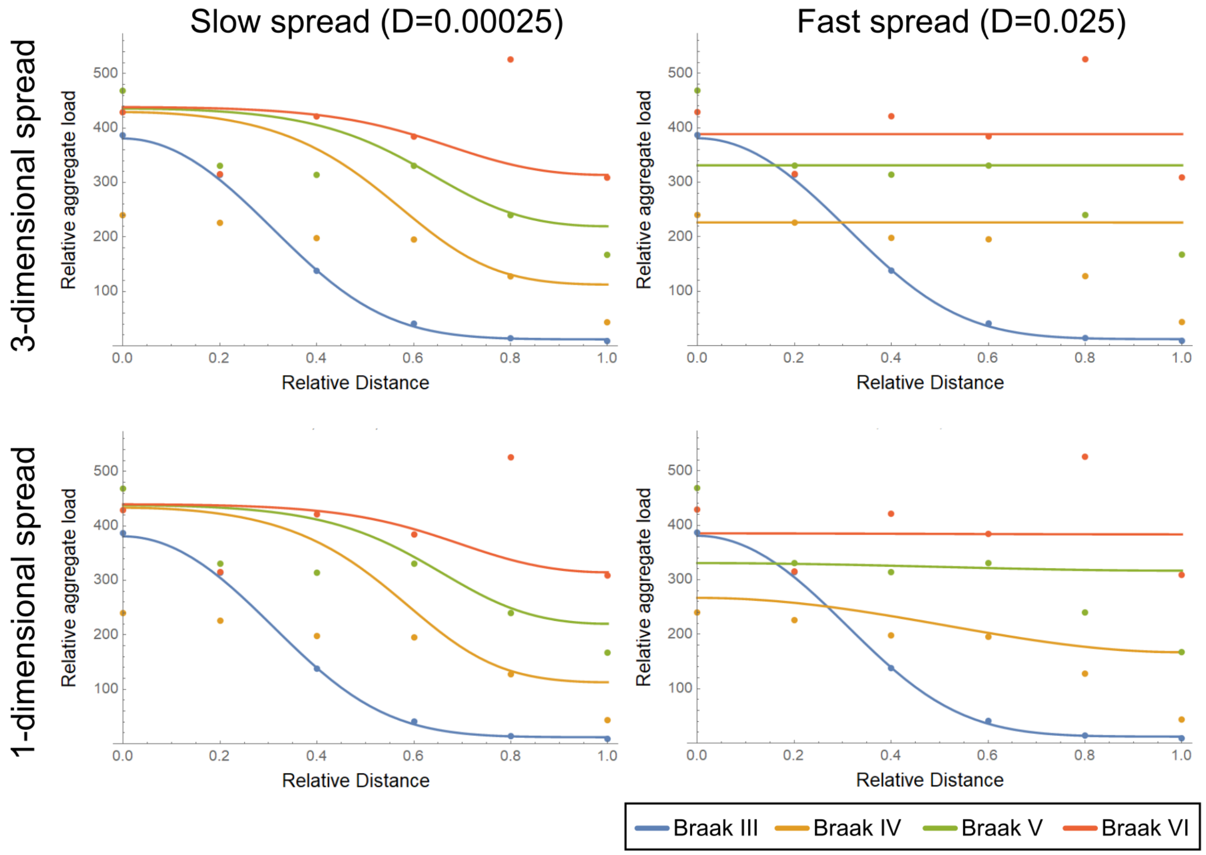


Figure S10: Comparison of 3-dimensional and 1-dimensional match to experimental data, from numerical integration of equations S2 and S3. According to the fits of the data, we chose $\kappa = 0.15$ years$^{-1}$. Temporal separation between simulations of different Braak stages are fixed by the Braak staging data. Effective diffusion coefficients are also fixed, thus only the time of switch between Braak stage III and IV is adjusted to match the data.

#### 2.4 Dissecting multiplication and growth

It can be shown that very generally, for a growth-multiplication type mechanism, the overall rate of increase in aggregates is given by

$$\kappa = \sqrt{k_{\text{growth}} k_{\text{mult}}} \tag{S12}$$

where $k_{\text{growth}}$ and $k_{\text{mult}}$ are the rates of growth and multiplication, respectively. Moreover, the average size of the aggregates, $\mu$, in number of monomeric units, is given by

$$\mu = \sqrt{\frac{k_{\text{growth}}}{k_{\text{mult}}}} \tag{S13}$$

Therefore, given the overall rate, $\kappa$, and the average size $\mu$, one can determine the rates of growth and multiplication by

$$k_{\text{growth}} = \mu\kappa \qquad k_{\text{mult}} = \frac{\kappa}{\mu} \tag{S14}$$

Jackson et al.[34] measured an average length of tau fibrils of 176 nm in P301S mice. Given a beta-sheet separation of 0.47 nm and assuming there are two tau monomers per layer of the structure[35], 176 nm corresponds to 750 monomers. To account for the fact that Jackson et al.[34] measured these sizes in P301S mice while Fitzpatrick et al [35] analysed fibrils form AD, we give a conservative range of likely $\mu$ to be between 100 and 10000 tau monomers, with the likely size on the order of 1000 monomers. We use the same range for both P301S mice and AD.

### 3 Relation to *in vitro* models

Due to the wealth of data and high level of control over experimental conditions, such as the monomer concentrations, significantly more detailed mechanistic information can be extracted from measuring the aggregation of a purified protein *in vitro*. In particular, we are usually able to determine the detailed mechanism of aggregate multiplication. The in vivo data analysed here do not permit this level of detail to be extracted, however, the more detailed models we developed in the context of in vitro aggregation can be approximately mapped to the coarse-grained model used here. While the details of how all processes and their various extensions map onto the coarse-grained models used her is beyond the scope of this work, we give here some general guidelines on how the 2 can be related. Spatial inhomogeneities are not generally considered in vitro, so we here simply compare the spatially independent version of equation S2 to the in vitro models. After some simplifications, the models of in vitro aggregation take the form of following non-linear coupled differential equations:

$$\frac{dP}{dt} = k_n m(t)^{n_c} + k_2 m(t)^{n_2} M(t) \tag{S15}$$

$$\frac{dM}{dt} = 2k_+ m(t) P(t) \tag{S16}$$

where $M(t)$ and $P(t)$ are the first and zeroeth moments of the fibril size distribution, i.e. the fibril mass and number concentrations, respectively and $m(t)$ is the free monomer concentration. The other parameters denote rate constants and reaction orders of the different processes on the pathway to aggregate formation. While the solutions of these equations generally take more complex forms than equation S4 derived here, they are generally similar in form, i.e. sigmoidal with an initially exponential increase, followed by a plateau region. *In vitro*, this plateau emerges from imposing conservation of mass via $M(t)+m(t) = m_{\text{tot}}$, *in vivo* it is an experimental observation and may arise due to a number of reasons, such as a decrease of protein synthesis in regions with high aggregate concentrations. For relating the models, this plateau region is of little importance and we instead focus on the early time exponential increase. A standard early time approximation *in vitro* is to set $m(t) = m_0$, as initially the depletion of monomer is negligible. This results in the following linear differential equations

$$\frac{dP}{dt} = k_n m_0^{n_c} + k_2 m_0^{n_2} M(t) \qquad \text{(S17)}$$

$$\frac{dM}{dt} = 2k_+ m_0 P(t) \qquad \text{(S18)}$$

Ignoring the contribution from the spontaneous formation of aggregates directly from monomer (the first term in the equation for $dP/dt$), the solutions are of the form

$$P(t) = P_0 \exp\left[\sqrt{2k_+ m_0 k_2 m_0^{n_2}}\, t\right] \qquad \text{(S19)}$$

Note that the exponential rate, $\sqrt{2k_+ m_0 k_2 m_0^{n_2}}$ is the geometric mean of the rate of elongation, $2k_+ m_0$, and the rate of multiplication, $k_2 m_0^{n_2}$.

The early time solution for the equation we use here, i.e.

$$\frac{dP}{dt} = \kappa P(t)(1 - P(t)) \approx \kappa P(t) \qquad \text{(S20)}$$

where $\kappa$ is the replication rate, is given by

$$P(t) = P_0 \exp[\kappa t] \qquad \text{(S21)}$$

We therefore identify $\sqrt{2k_+ m_0 k_2 m_0^{n_2}} \rightarrow \kappa$. In other words the replication rate here coarse-grained the processes of growth and multiplication into one rate and also subsumes their reaction orders and rate constants into one term. We can to some extent still dissect this replication rate by making use of the fact that growth and multiplication affect the average length differently, as we have done in the main part of this work, but the monomer dependence of processes is not accessible. Given the fact that many additional processes will be present *in vivo* that affect both multiplication and growth, the rates extracted from in vivo data represent effective rates of growth and multiplication rather than exact equivalents of their *in vitro* counterparts. However, to what extent growth or multiplication are affected points towards how these additional processes present *in vivo* affect aggregation.

[3] Georg Meisl, Timothy Kurt, Itzel Condado-Morales, Cyrus Bett, Silvia Sorce, Mario Nuvolone, Thomas C. T. Michaels, Daniel Heinzer, Merve Avar, Samuel I. A. Cohen, Simone Horneman, Adriano Aguzzi, Christopher M. Dobson, Christina J. Sigurdson, and Tuomas P. J. Knowles. Scaling analysis reveals the molecular mechanism and rates of prion replication in vivo. *manuscript under preparation*, 2019.
